# Social Learning of a Spatial Task by Observation Alone

**DOI:** 10.1101/2022.03.28.485797

**Authors:** Thomas Doublet, Mona Nosrati, Clifford G. Kentros

## Abstract

Interactions between conspecifics are central to the acquisition of useful memories in the real world. Observational learning, i.e., learning a task by observing the success or failure of others, has been reported in many species, including rodents. However, previous work in rats with NMDA-receptor blockade has shown that even extensive observation of an unexplored space through a clear barrier is not sufficient to generate a stable hippocampal representation of that space. This raises the question of whether rats can learn a spatial task in a purely observed space from watching a conspecific, and if so, does this somehow stabilize their hippocampal representation? To address these questions, we designed an observational spatial task in a two-part environment that is nearly identical to that of the aforementioned electrophysiological study, in which an observer rat watches a demonstrator animal to learn the location of a hidden reward. Our results demonstrate that rats do not need to physically explore an environment to learn a reward location, provided a conspecific demonstrates where it is. We also show that the behavioral memory is not affected by NMDA receptor blockade, suggesting that the spatial representation underlying the behavior has been consolidated by observation alone.

## INTRODUCTION

In humans and many animals, new behaviors may be learned through the observation of a conspecific’s experience. Observational learning has been reported in invertebrates (Worden and Papaj, 2005), vertebrates such as birds and fish (Dawson and Foss, 1965) (Laland and Williams, 1998), mammals (Bunch and Zentall, 1980) and humans (Bandura, Ross and Ross, 1961).

Rodents can adjust their behavior to the behavior of conspecifics using visual information (Worden and Papaj, 2005) (Keum and Shin, 2019). By observing a conspecific, rodents can more quickly learn complex tasks such as pressing a lever to obtain rewards or cooperative behavior in social games (Zentall and Levine, 1972) (Heyes and Dawson, 1990) (Viana et al., 2010). Interestingly, observing a conspecific’s failure to succeed is more informative for learning a task through observation than observing its success (Templeton, 1998).

All known studies on observational learning of a spatial task imply the learning of efficient strategies to accomplish the task or include subjects with previous self-experience of that space (Leggio et al., 2000) (Leggio et al., 2003) (Petrosini et al., 2003) (Takano et al., 2017) (Bem et al., 2018). Leggio demonstrated the role of the cerebellum in learning successful strategies from conspecific experience in various spatial tasks (Morris water mazes). Takano claimed that rats can learn efficient strategies for success in a spatial task from inefficient experiences of conspecifics navigating in a known space. Finally, Bem showed that observing a conspecific lead to more relevant search strategies. Furthermore, Bem showed that observing an experienced demonstrator is beneficial only when what is observed is relevant or novel enough to complement existing knowledge. Unfortunately, none of these studies indicate whether it is possible to develop a stable representation of an observed, unexplored space.

Rodents can independently remember locations in a radial arm maze (Olton, 1977) or find a hidden platform in a water maze (Morris, 1984). Tolman theorized that animals may have an internal spatial map that could represent geometric coordinates of the environment and effectively aid navigation even when visiting a space for the first time (Tolman et al., 1946) (Tolman, 1948). The spatial firing fields of the hippocampus and associated cortices has been proposed to be the neural instantiation of the cognitive map of space theory (Fyhn et al., 2004) (Buzsáki and Moser, 2013) (Moser, Moser and McNaughton, 2017).

These spatial firing fields include place cells (O’Keefe and Dostrovsky, 1971) (O’Keefe and Nadel, 1978) (Wilson and McNaughton, 1993), grid cells (Hafting et al., 2005) (Sargolini et al., 2006) (Barry et al., 2007), border cells (Solstad et al., 2008) (Savelli, Yoganarasimha and Knierim, 2008), and head-direction cells (Ranck, 1985) (Taube, Muller and Ranck, 1990). Place cells, for example, are hippocampal neurons that are selectively activated when an animal occupies a particular location of a particular environment, referred to as its place field. The processes that control the generation of a hippocampal representation of an environment remain poorly understood, including whether they can be formed in spaces that are simply observed or whether direct experience of the space is necessary. The difficulty with this is that one cannot know that a cell has a place field at a particular location until the animal visits that location.

However, the only electrophysiological study to directly examine whether rats can create a stable place cell map of an unexplored space found the opposite (Rowland, Yanovich and Kentros, 2011). Rats were trained in 2 concentric boxes, with the inner box made of clear plexiglass and the outer box containing the only available cues. During observational training in the inner box, they could see the outer box but could not physically explore it. On the test day, the animals were able to explore the entire environment either with or without NMDA receptor blockade, which prevents stabilization of a newly formed place cell map but does not destabilize a previously formed one (Kentros et al., 1998). This allowed them to show quite clearly that the map was stabilized only after direct exploration (i.e., the place fields of the drug animals were stable in the inner box but unstable in the outer box, while the saline ones were stable everywhere).

However, this raises the question as to whether a rat *cannot* learn spatial information purely by observation, or whether they simply had no reason to do so. We therefore modified this maze by adding 12 pebble-covered food wells to the outer box, one of which contained a hidden reward. The animal in the inner box had to learn the goal location purely by observation of a trained conspecific’s behavior in the inaccessible outer box. Thus, this novel observational learning task combines both spatial and social learning in one.

## MATERIALS AND METHODS

### Animals

Animals were bred locally at NTNU. They were kept in a 12 h LD light cycle and fed ad libitum. They were housed in environmentally enriched cages in a humidity and temperature-controlled environment. 45 male Long Evans rats were included in the present study (3-7 months old at the time of testing). All procedures took place during the light cycle.

All procedures were approved by the National Animal Research Authority of Norway. They were performed in accordance with the Norwegian Animal Welfare Act and the European Guidelines for the Care and Use of Laboratory Animals (directive 2010/63/UE).

### Experimental Design

We tried to keep the experimental design as similar to that previously reported with place cell recordings (Rowland, Yanovich, and Kentros, 2011), only adding the social transmission of the spatial task. Thus, experiments were conducted in a customized behavioral apparatus that consisted of two square boxes: a transparent Plexiglas inner box (50 × 50 cm) within an opaque outer box (100 × 100 cm) with asymmetric spatial cues available to the animal. Additionally, twelve symmetrically distributed wells were included in the outer space between the two boxes. An equal number of pebbles covered each well to hide the potential reward (chocolate loops, Nestle). Before each animal was introduced into the apparatus, the pebbles that had a cue were replaced with new ones. An accessible but not visible reward was placed in one of the wells. Rewards were also placed evenly under the entire perforated floor of the apparatus to ensure a uniform odor in all wells and to minimize the possibility that a rat could identify the correct well by odor. The reward had an 8.3% probability of being found by the rats by chance.

### Behavioral Testing

All rats were familiarized to the experimental environment daily for at least three sessions of thirty minutes each. During this time, the rats were confined in the transparent inner box, which was located within the outer box (as shown in **Figure 1**). This allowed the inner box to be experienced directly, while the outer box could only be observed. At the end of each familiarization session, the rat was returned to its home cage for at least 8 hours. The floor, pebbles, and walls of the maze were cleaned with 90% ethanol after each session. Animals were habituated to the reward in their home cage daily before the start of the experiment.

**FIGURE 1.**
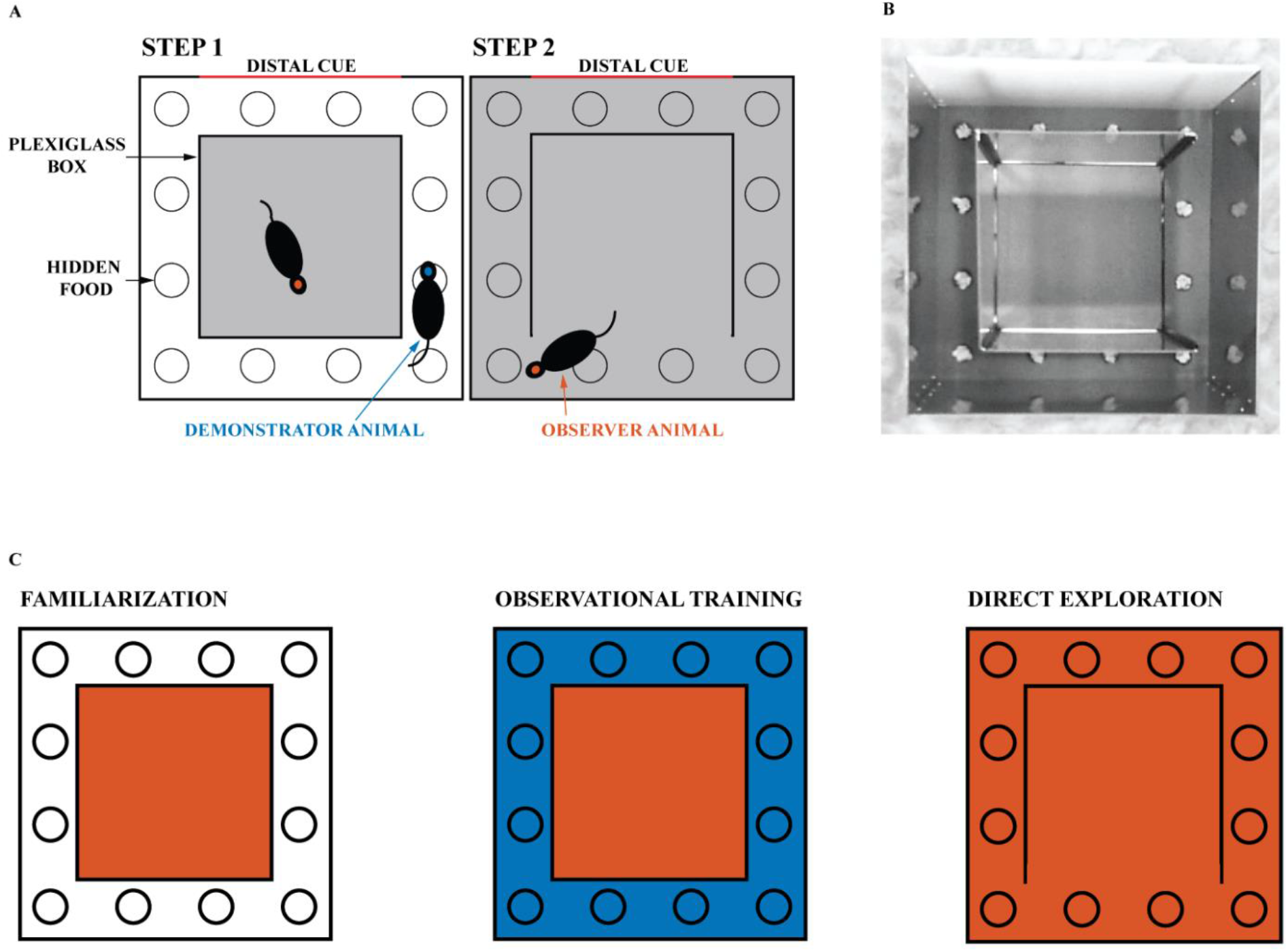
Experimental design. (A) The experimental environment consisted of a transparent inner box and an opaque outer box. The gray areas indicate the regions explored by the tested rat. (B) Image of the experimental apparatus with the right wall of the transparent inner box open. The reward is hidden in one of the 12 wells and covered with gravels. One of the four walls of the opaque outer box is white and provides a distal cue to the animals. (C) Schematic representation of the experiment. The familiarization phase, in which the experimental animal is confined to the inner box, is followed by the observational training phase, in which it can observe the demonstrator animal navigating the outer space (blue). Finally, on the day of direct exploration, the observer animal is allowed to navigate in the observed space. One session is held daily, for a total of 9 sessions (3 for familiarization, 5 for observational training, and 1 for direct exploration). The red and blue areas correspond respectively to the space that the observer and demonstrator animals can physically explore.

Rats were tested for task success (i.e., number of erroneous attempts) and time taken to find the reward (i.e., latency) during their first direct exploration of the outside space. Subjects were divided into naive (n=27) and trained animals (n=18). Naive animals were tested for the ability to find the reward without any observational training. After at least twenty consecutive successful trials, the naive animals became **demonstrator** animals (see **Figure Supp. 1**). **Observer** animals were trained on the location of the reward by the demonstrator animals. During training sessions, each observer animal was paired with the same demonstrator animal, and the reward was always in the same single well (see **Figure 1A-B**). Observational training consisted of five rewards (for the demonstrator) daily for five consecutive days (see **Figure 1C**). Each new reward was made available five minutes after the previous reward was discovered. Animals were not removed during rebaiting to avoid stress and disengagement on the task (Cloutier, 2015). Instead, all wells were manipulated with obscured vision for the animals. Observational training was completed after 25 rewards were found by the demonstrator animal in the presence of the paired observer located in the plexiglass inner box. After observational training was completed, the observer rat was allowed to explore the outside space and find the reward itself. As in our previous study (Rowland, Yanovich, and Kentros, 2011), the outside space was entered through the opening of a plexiglass wall opposite the reward well. The reward well was in a different location for each pair of animals to mix up the cues. To increase social interaction, the animal pairs were siblings housed in adjacent home cages. Finally, the NMDA receptor antagonist CPP [(±)- 3-(2-carboxypiperazin-4-yl) propyl-1-phosphonic acid, 10 mg/kg, Sigma] was injected intraperitoneally in a subset of 5 observer animals before the first direct exploration of the outside space (but after the observation stage was complete).

Success and latency of observer and naive groups were compared. A trial was considered successful if the animal made no mistakes prior to digging in the correct well. A mistake was counted as active digging in an unrewarded well. Pebble removal that was not performed with the head or front limbs or while the animal was running was not counted as active digging. Evaluation of animal performance by experimenters was confirmed by video analysis of two blinded students independent of the study who reached identical conclusions (2 students quantified trials of 10 animals). A separate cohort of observer animals was tested with no reward present during the initial outside direct exploration (see **Figure 1C**). A third cohort of observer animals was tested one hour after CPP injection, with no reward present during the first outside direct exploration (see **Figure 1C**).

### Data Analysis

All data were analyzed using the average time taken to find the reward from entering the outside space, the total number of mistakes made, and the percentage of successful animals for each trial. All values were expressed as mean ± standard error of the mean (SEM). All behavioral data were analyzed using the Pearson chi square test and the unpaired mean difference between control and test, as indicated, using SPSS software (IBM) and MATLAB (Ho, 2019). All tests were two-tailed tests. For the unpaired mean difference between control and test, 5000 bootstrap samples were taken, and the confidence interval is bias-corrected and accelerated. Reported P values are the likelihoods of observing the effect size if the null hypothesis of zero difference is true. Effect sizes and confidence intervals (CI) are reported as: Effect size [CI width lower bound; upper bound].

Cohort and sample sizes were reported in the text and figures. Statistical significance was set at p < 0.05 ‘‘∗’’, p < 0.01 ‘‘∗∗’’ and p < 0.001 ‘‘∗∗∗’’.

## RESULTS

### Experiment 1: Learning a reward location in naive rats

Previous studies in rodents have found that learning a spatial task follows a logarithmic curve of success until a plateau is reached. Our task described in **Figure 1** followed the same rule. **Figure 2A** shows the progression of success for a naive animal in this task. A success is counted if the animal found the reward on the first try without digging in other wells. The probability of finding the reward was 8.3% (1 well out of 12). The probability of success on the first reward for naive animals is comparable to chance (12.5%). The percentage of successful naive animals at the first 15 rewards were, respectively: (1) 12.5% ± 8.5 (mean percentage ± SEM); (2) 55.6% ± 12.1; (3) 83.3% ± 9.0; (4) 81.3% ± 10.1; (5) 92.9% ± 7.1; (6) 89.5% ± 7.2; (7) 94.7% ± 5.3; (8) 100%; (9) 100%; (10) 100%; (11) 94.7% ± 5.3; (12) 100%; (13) 100%; (14) 100% and (15) 100% (n=14). Success at the first direct exploration was statistically different from the second (Pearson chi-square = 6.88, 99.9% confidence, n_1_= 16 and n_2_= 18). Similarly, success at the second direct exploration was statistically different compared to the third (Pearson chi-square = 3.27, 95% confidence, n_2_= 18 and n_3_= 18).

**FIGURE 2.**
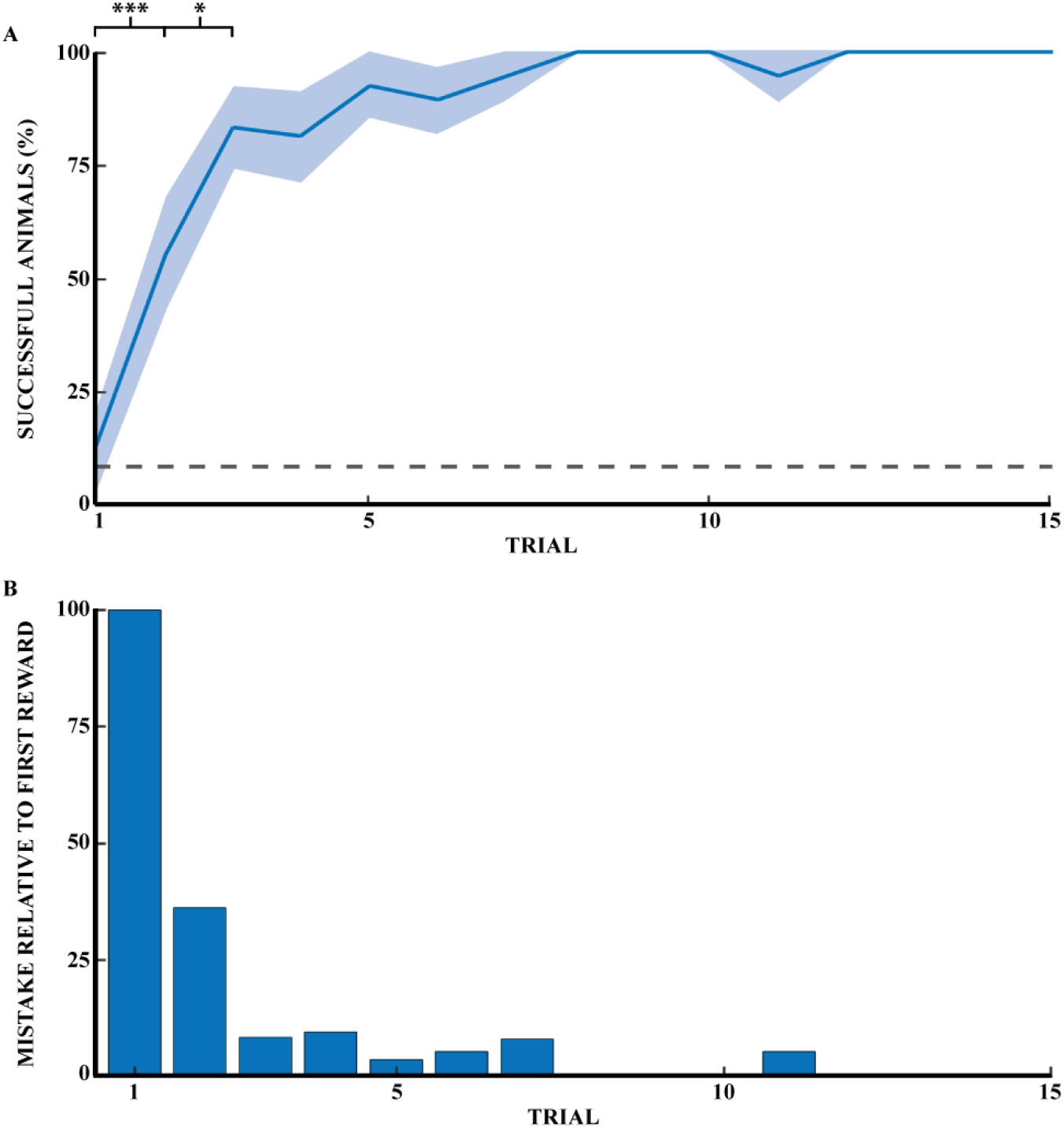
Spatial memory task learned through exploratory experience. **(A)** Learning progress of naive rats across 15 reward retrievals (3 days) calculated as percentage of successful animals for each trial (n= 14). Error bars are mean ± standard error of the mean (SEM). Gray dashed line represents success by chance. **(B)** Number of mistakes per trial by naive rats across 15 reward retrievals (n= 14). Number of mistakes is the average normalized number of mistakes made for each reward, relative to the first trial. * p < 0.05, *** p < 0.001.

**Figure 2B** shows the reduction of mistakes across 15 reward retrievals. A mistake was counted as actively digging in a non-target well, with a maximum number of mistakes per trial of 11. This figure shows that naive animals stopped making errors after 11 trials (n=14 rats). Mistakes are shown here relative to the first direct exploration. Animals were monitored until 20 consecutive successes, but only the first fifteen rewards were shown in **Figure 2**. Recall that a naive rat was considered a demonstrator rat after at least 20 consecutive successful trials, and thus the observer rats were effectively exposed to the perfect performance of the task by the demonstrator animal.

Finally, the time it took the naive animals to find each reward (**Figure 3B**, blue curve) decreased similarly from the first reward and reached a plateau after 4 rewards. The time taken by naive animals to find each of the first five rewards was: (1) 1515.6 ± 484.4; (2) 277.3 ± 89.5; (3) 347.6 ± 189.3; (4) 64.9 ± 15.5 and (5) 110.9 ± 30.7 seconds (n= 17).

From this we can conclude that the task needs experience to be completed and cannot be achieved without it.

### Experiment 2: Learning the location of a reward through social observation

To investigate whether learning the location of a hidden reward is possible through social observational training, we trained observer rats to find the location of a hidden reward using demonstrator animals (5 trials daily for 5 consecutive days). We then had the observer animals go out to explore the observed space and find the reward (see **Figure 1**). The observer group successfully found the reward in 100% of the animals without error during their first direct exploration of the outside space (N=6) (**Figure 3A**). All subsequent direct explorations were also 100% successful (n=15 trials, 5 animals). Performance on the first direct exploration was statistically different from that of the naive animals (Pearson chi-square = 14.44, 99.9% confidence, n_n_= 16 and n_o_= 6). Performance across trials did not differ significantly between observer animals.

**FIGURE 3.**
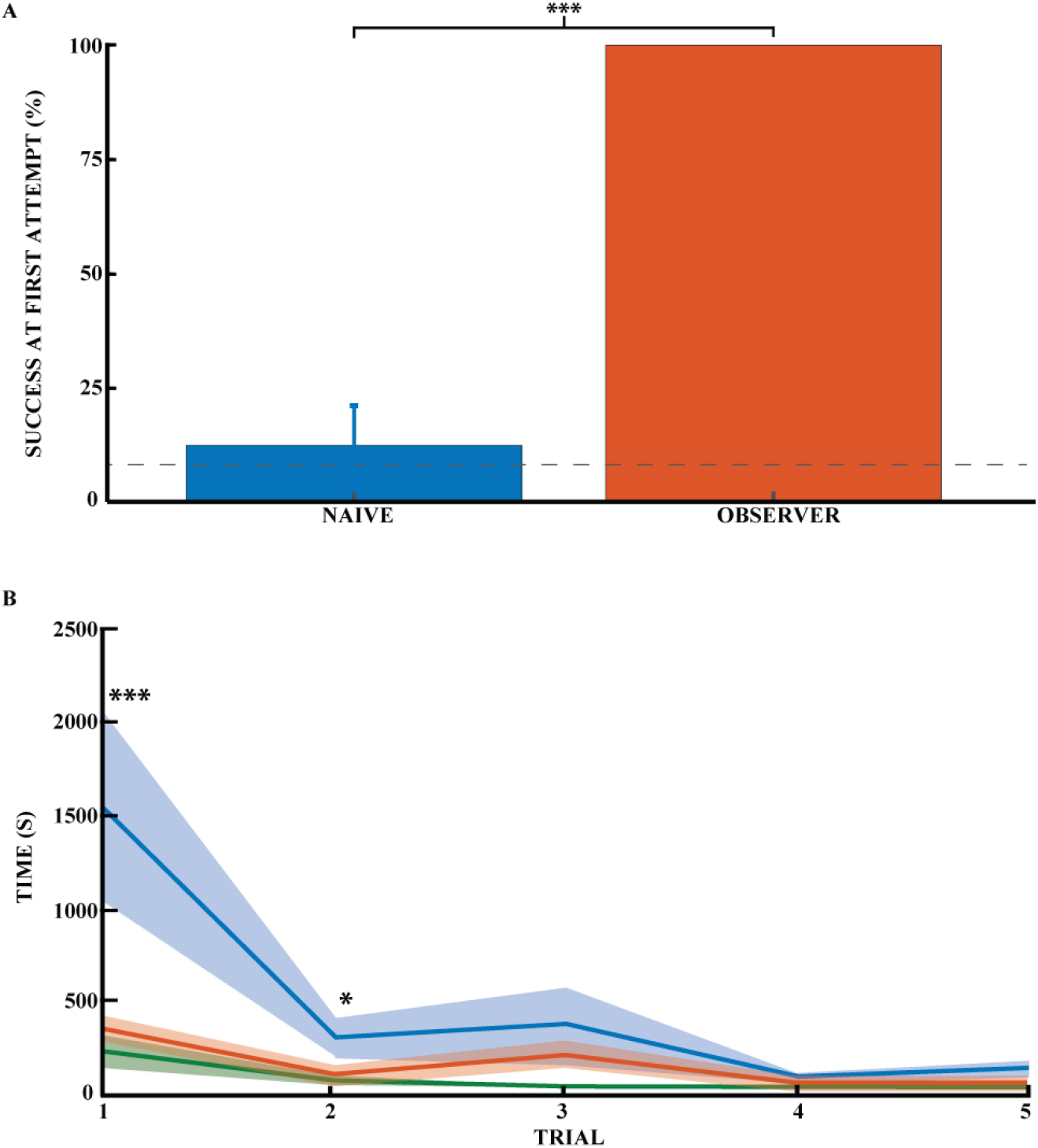
Spatial memory task learned by observational experience in an unexplored environment. (A) Effect of learning an unexplored space by observation by percentage of success on the task for naive (blue) and observer animals (red) on the first direct exploration. Performance on the first direct exploration was statistically different for the observer animals compared to the naive animals (Pearson chi square= 14.44, 99.9% confidence, n naive = 16, n observer = 6). Error bars are mean ± standard error of the mean (SEM). Gray dashed line represents success by chance. (B) Effect of learning the unexplored space by observation using the average time to find the reward across trials (n naive = 17, n observer = 5). Performance on the first and second direct explorations was statistically different in observer (red) compared with naive animals (blue) (unpaired mean difference on first reward = -1.17*10^3^, 99.9% confidence; unpaired mean difference on second reward = -1.85*10^2^, 95.0% confidence). Demonstrator (green) for comparison. Error bars are mean ± standard error of the mean (SEM). * p < 0.05, *** p < 0.001.

While latency towards reward is a common measure of spatial performance, it is not particularly informative in this case because the animals invariably first explore the novel space prior to engaging with the spatial task. Still, there was an appreciable difference between trained and untrained animals. The animals in the observer group required much less time to find the first rewards (**Figure 3B**, red curve). It was around half the time it took for naïve animals (**Figure 3B**, blue curve). Thus, time to reward was significantly different between the naive and observer groups for the first two rewards. The unpaired mean difference between naive and observer animals was -1.17_*_10^3^ [99.9% CI -2.31_*_10^3^, -4.12_*_10^2^] for the first trial and -1.85_*_10^2^ [95.0% CI -3.85_*_10^2^, -20.1] for the second trial. The latency of observers was not significantly more than for demonstrators (**Figure 3B**, green curve), The unpaired mean difference between observer and demonstrator animals was 1.44e+02 [95.0% CI -1.17_*_10^2^, 2.85_*_10^2^] for the first trial and 49.0 [95.0% CI - 18.1, 1.66_*_10^2^] for the second trial. So far as errors go, the observer and demonstrator groups performed comparably even during the first two trials (1) -1.44_*_10^2^ [95.0% CI -2.87_*_10^2^, 1.09_*_10^2^] and (2) 49.0 [95.0% CI -16.5, 1.67_*_10^2^]. The time it took the naive and demonstrator rat groups to obtain the rewards was significantly different for all first five rewards (1) -1.32_*_10^3^ [99.9% CI - 2.65_*_10^3^, -5.99_*_10^2^]; (2) -3.04_*_10^2^ [99.9% CI - 8.6_*_10^2^, -35.4]; (3) -3.37_*_10^2^ [99.9% CI -1.08_*_10^3^, -37.8]; (4) -56.6 [99.9% CI -1.17_*_10^2^, -19.4] and (5) -1.04_*_10^2^ [99.9% CI -2.35_*_10^2^, -46.6]).

Thus, unlike the naive animals, the observer and demonstrator groups did not make mistakes in accomplishing the task. In addition, the time it took the observer animals to successfully complete the task was comparable to that of the demonstrators, but both groups were statistically faster than the naive animals. Observer animals tend to explore the maze once or twice before engaging in the task. The time required to learn and successfully complete the task is coherent with the literature for such a naturalistic social learning task (no food deprivation, no time limit). This task is very time consuming, and the latency required for the animals to find the reward makes time less meaningful than success or failure in the task.

We controlled for cleaning quality to ensure that odor was not a factor for animals to navigate to the reward via olfaction. When two naive rats explored the outside area for the first time within 30 minutes, the first well dug by the second animal was compared to the reward location of the previous animal. Among the 12 pairs of animals, the second rat never dug the first animal’s reward well first. This result confirmed that cleaning within two sessions was effective and had no undesirable effect on the outcome of the next animal.

### Experiment 3: Is the behavior dependent on olfactory cues?

Even though reward odor was distributed throughout the maze, it is possible that the rats were still capable of using olfactory gradients to solve the task without observational spatial learning. To investigate the influence of reward odor on animal navigation, we compared the ability of naive animals to dig in the correct well with and without reward. **Figure 4A** shows the average number of mistakes on the first trial (how many incorrect wells were dug before the correct one) for the rewarded and non-rewarded naive animals. For the latter animals, no accessible reward was hidden, so we can rule out navigation by smell to the correct well. Thus, for this group, the number of mistakes made before digging in a given well would be completely random, so we can control for whether the smell of the hidden chocolate loop might provide a cue to reduce the number of mistakes made. The difference between the two naive groups was significant, indicating that the reward odor could reduce the number of errors made by the animals in the rewarded condition (Pearson chi square = 15.44, 95% confidence, n_R_ = 16 and n_NR_= 7). The number of mistakes made in the first exposure was 4.4 (SEM = 0.8) for unrewarded naive animals and 2.0 (SEM = 0.3) for rewarded animals. However, the number of successful animals appeared to be independent of the presence of a reward for naive animals. Both groups were close to chance (8.3%) at the first direct exploration with 12.5% (n_R_ =16, SEM = 8.5) and 0% (n_NR_=7) for rewarded and non-rewarded animals, respectively (**Figure 3A** for naive rewarded and **Figure 4B** for naive non rewarded). The difference between the two naive groups was not statistically significant (Pearson chi square = 0.98, n_R_= 16, and n_NR_= 7). The time required to find the reward at first exposure was also not significantly different, 1515.6 ± 484 and 1404 ± 744 seconds, respectively (unpaired mean difference is -1.12e+02 [95% CI - 2.04e+03, 3.25e+03]).

**FIGURE 4.**
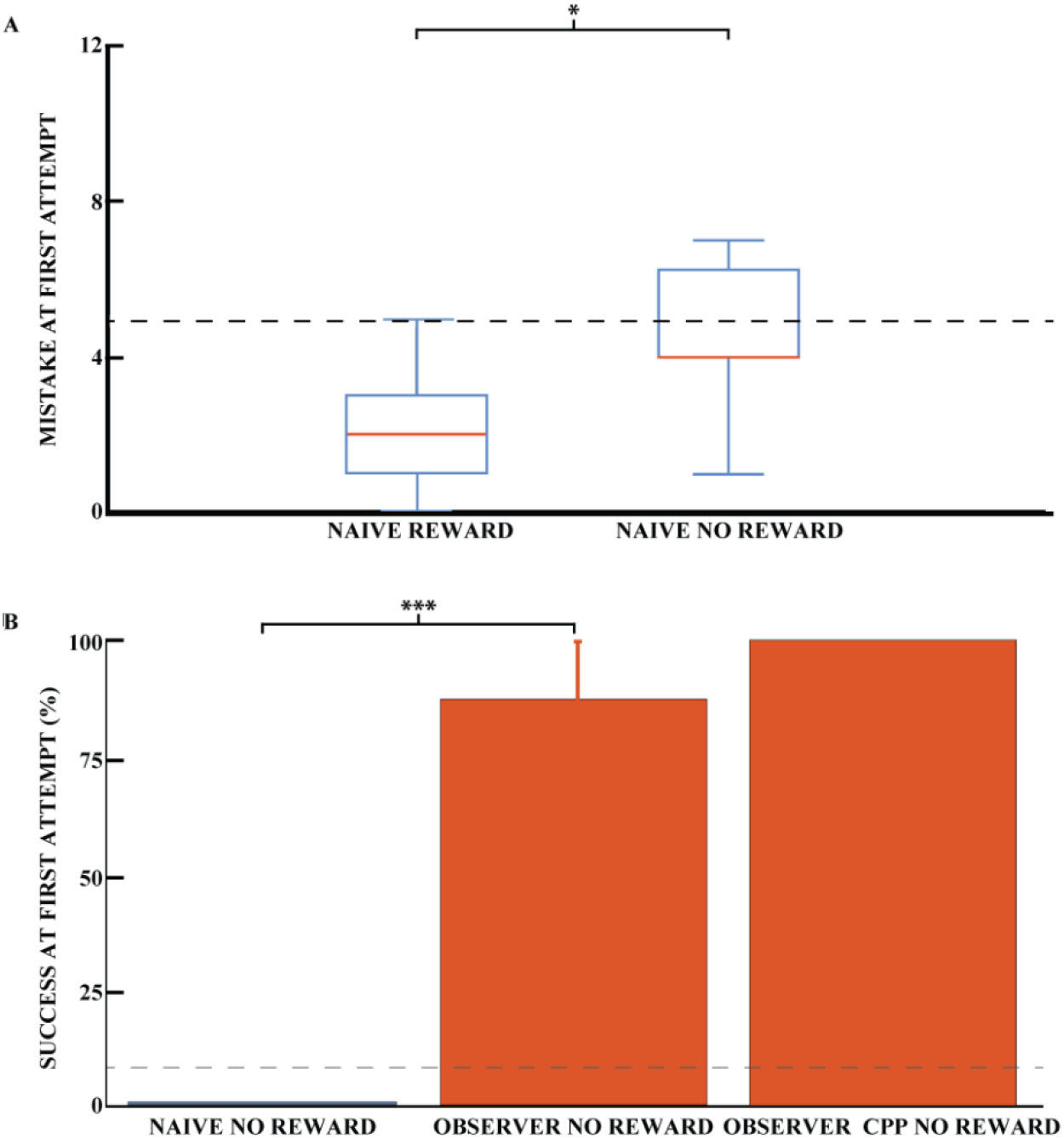
Success on the spatial task is independent of olfactory cues. (A) Mean number of mistakes on the first trial for rewarded and unrewarded naive animals. Performance on the first direct exploration was statistically different for rewarded and non-rewarded naive animals (Pearson chi-square= 15.44, 95% confidence, n naive rewarded = 16, n naive non-rewarded = 7). Error bars are mean ± standard error of the mean (SEM). Gray dashed line represents success by chance. (B) Effect of learning an unexplored space by observation using the percentage of success in the unrewarded task for naive (blue) and observer animals (red) on the first direct exploration. Performance on the first direct exploration was statistically different for observer animals without reward (red) compared to naive animals without reward (blue) (Pearson chi-square= 10.50, 99.9% confidence, n naive animals without reward = 7, n observer without reward = 8). No statistical difference was found between unrewarded observer animal control and CPP groups (n observer non-rewarded = 8, n observer non-rewarded CPP = 5). Error bars are mean ± standard error of the mean (SEM). Gray dashed line represents success by chance. * p < 0.05, *** p < 0.001.

To preclude localization of the reward by the sense of smell of the observer animals, the reward was removed after observational training but before the first outside direct exploration for a cohort of observer animals. Each of these observer animals was trained with a paired demonstrator that performed 25 trials, similar to previously described. Observer animals that explored the outside environment without reward after observational training were 87.5% successful on their first direct exploration (n= 8 animals, SEM = 12.5), see **Figure 4B**. Only one observer animal made an error in the task, and he made 6 mistakes during his first direct exploration. The percentage of success on the first trial was not statistically different between the rewarded and non-rewarded observer cohorts (Pearson square= 0.81, n_R_ = 6 and n_NR_= 8, respectively), nor was the number of mistakes (unpaired mean difference is 0.75 [95% CI 0.0, 3.75]). The difference in mistakes between the unrewarded naive and observer groups was statistically significant, as was the difference in mistakes between the rewarded groups (Pearson square= 10.50, 99.9% confidence, n= 7 and n= 8, respectively). Rewarded and unrewarded observer animals showed similar performance, ruling out a possible olfactory influence on task success.

### Experiment 4: A stable representation of space is formed before the first direct exploration

To confirm that a stable representation of space can be formed before the first physical direct exploration of a space, we injected CPP (an NMDA receptor antagonist). CPP prevents stabilization of a newly formed hippocampal representation of an environment but does not destabilize an already formed one (Kentros et al., 1998). Interestingly, observer animals that explored the observed space one hour after an injection of the NMDA receptor antagonist CPP performed similarly to animals that did not receive an injection (**Figure 4B**).

These observer animals with CPP that explored the outside environment without reward were 100% successful on their first direct exploration (n= 5 animals). The three observer cohorts (observer, observer unrewarded and observer unrewarded with CPP) share comparable chances of success in the task.

During these unrewarded experiments (Experiments 3 and 4), the animals performed the task only once because of extinction of the memory.

For all animals, the percentage of success on the first trial was statistically different when the naive and observer groups were compared (Pearson chi square= 23.25, 99.9% confidence, n= 23 and n= 19, respectively). The percentage of success on the first trial was 8.7% (SEM = 6.0) for naive animals and 92.3% (SEM =7.7) for observer animals, clearly indicating knowledge of the goal location from observation alone

## DISCUSSION

The behavioral studies presented are to our knowledge the first to directly investigate the performance of rodents in a spatial task in an unexplored space with training exclusively based on observation of a conspecific performing that task. We found that this observation led to highly significant improvements in both accuracy and latency towards the goal as compared to naïve animals, even though the structure and operant nature of the task means that the observer animals’ native tendency to explore a novel space (the outer box) competes with their engagement with the digging task.

The performance improvement followed a learning curve similar to that described in classical learning theory (Wright, 1936) (Anzanello and Fogliatto, 2011). In this model, performance on a repetitive task improves through repetition. A learning period is then followed by a learned period in which performance reaches a plateau. **Figure 2** shows the success rate of naive animals in the task for each trial. We can then track performance in the task as experience increases. The percentage of successful animals increases significantly from reward one to reward two and from reward two to reward three and so on.

**Figure 3** compares the success rate (digging in the right well) in the first trial for naïve versus observer animals. The observer group clearly outperforms the naive group of animals (100% success versus 12%; chance is 8.3%). The situation is similar for the second reward. Moreover, the same conclusion can be drawn for the time taken to find the reward in the first two trials. Furthermore, the observer animals did not make a mistake in the next thirteen trials and thus do not fit a learning curve.

These results imply that the observer animals learned the goal location by watching a conspecific, as they were able to find the reward successfully from the first trial. While certainly some of the performance difference between observers and naïve animals had to do with observing nonspatial features of the task (e.g. the fact there is a reward that you have to dig for), the goal location as well was learned by observation because 1) the observer animals outperformed the naïve animals from the first trial and not after several trials and 2) there is no improvement by additional exploratory learning in the observer animals, which contradicts previously described cases involving efficient strategies (Leggio et al., 2000) (Leggio et al., 2003) (Takano et al., 2017) (Bem et al., 2018). Comparison between rewarded and non-rewarded observer animals (**Figures 3 and 4**) shows no difference between the two cohorts in initial direct exploration of the observed space, ruling out the possibility that the animals’ sense of smell could help them navigate to the reward.

This suggests that animals trained by observation have a representation of the reward location before its first direct exploration. This is in sharp contrast to our previous study which clearly showed the opposite result: a stable hippocampal representation of a space required its direct experience (Rowland, Yanovich, and Kentros, 2011). The destabilization of the place fields in this task was caused by CPP injections as well, which have consistently destabilized newly formed place fields (Kentros et al., 1998) (Rowland, Yanovich, and Kentros, 2011) (Dupret et al., 2010) (O’Neill et al., 2010) but did not affect performance in this observational task. Since the only difference was the observational learning of a spatial goal location, this means that either the observed space was stabilized by observation alone, or that a stable place cell representation is not necessary for spatial task performance.

While these possibilities can only be disambiguated by electrophysiological recordings, the preponderance of evidence points to the first option. Bats and rats have a cognitive representation of a familiar space being explored by a conspecific (Omer et al., 2018) (Danjo, Toyoizumi and Fujisawa, 2018). In these two studies, the place cells of the observer animals fired relative to the position of the observed animal’s location, providing a neural basis for such a thing. Similarly, “preplay” suggest that rats can make a spatial representation from distance (Gupta et al., 2010) (Dragoi and Tonegawa, 2011) (Ólafsdóttir et al., 2015). The study most similar to this one showed that a trained demonstrator can only “teach” an observer animal if what is being observed is sufficiently relevant or novel (Bem et al., 2018). In their study, the observer had already physically experienced the observed space (thereby creating a stable place cell map of it) and just had to learn the location of the rewards in that space. Moreover, it is entirely consistent with the observation that increased attention to space increases the stability of a hippocampal representation (Kentros et al., 2004) (Muzzio et al., 2009). Remote (i.e., observational) exploration of a space may be far less capable of stabilizing its hippocampal representation (Rowland, Yanovich, and Kentros, 2011), but the rats in that study were given no reason to attend to the outer box. Perhaps if the animal pays enough attention to the space, it will stabilize its place cells of it.

Of course, the possibility that stable place cells are not necessary for spatial task performance cannot be ruled out since the present study has no electrophysiological recordings, but this would contradict most studies which have examined this idea. Transgenic animals with behavioral deficits in spatial tasks (Renaudineau et al., 2009) (Arbab, Pennartz, and Battaglia, 2018) (Rotenberg et al., 1996) tend to have unstable place fields, and a chemogenetic manipulation that led to hippocampal remapping led to clear deficits in spatial memory retrieval (Kanter et al., 2017). Still, it remains possible that “third-person” representations of space are formed distinct from more familiar forms of hippocampal spatial firing. Regardless, we have shown that rats can obtain sufficient knowledge of an unexplored space to successfully locate a hidden reward purely by observing a conspecific’s behavior. This task should therefore provide a means to explore both the structure of a cognitive map and the representation of a conspecific’s behavior.

## DATA AVAILABILITY

The original contributions presented in the study are included in the article/supplementary material, further inquiries can be directed to the corresponding author.

## AUTHOR CONTRIBUTIONS

TD and CK designed the study. TD and MN conducted the research. TD conducted statistical analyses. TD and CK wrote the manuscript. CK and TD did the project administration and supervision. All authors critically revised the manuscript and gave approval for publication.

## ACKNOWLEDGMENTS

We are grateful to Dr. David Rowland for his fruitful discussions, Drs. Bartul Mimica and Tuce Tombaz for their valuable discussions on the statistical analyzes in this manuscript. The Clawsons are thanked for their invaluable assistance in writing the manuscript. Finally, we thank all the members of the Kavli Institute and especially the animal technicians and veterinarians for their support and kindness.

