## Supplemental Figure 1 for "Social Learning of a Spatial Task by Observation Alone"

**Suppl. Figure1**

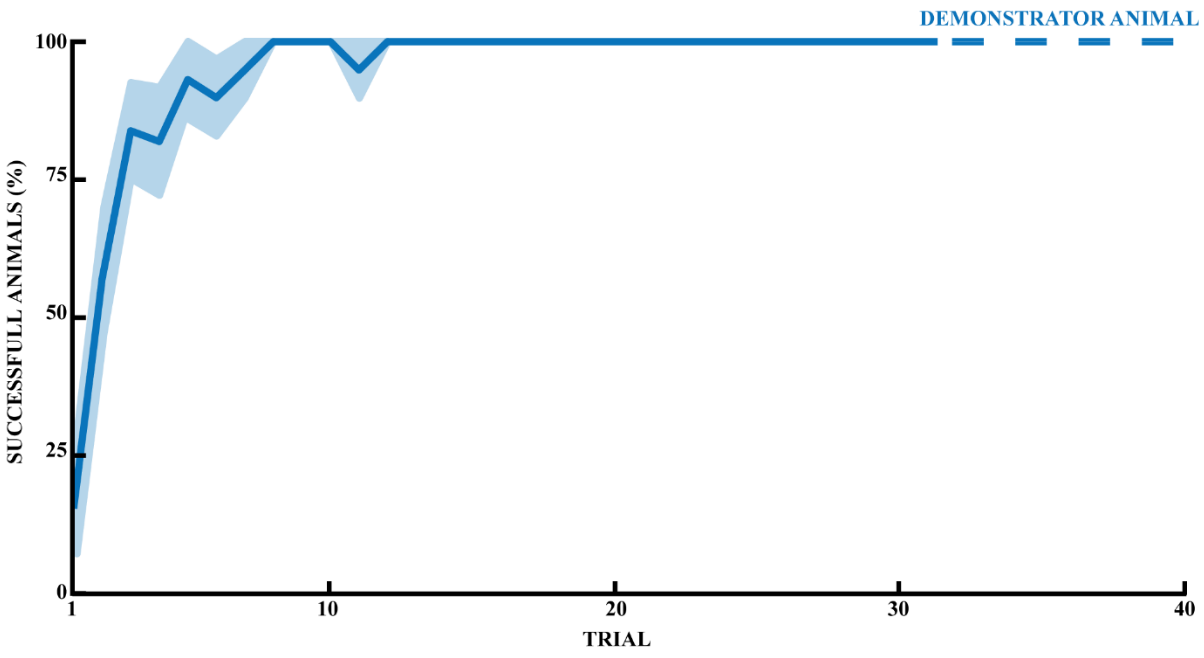


**FIGURE SUPP. 1** | Spatial memory task learned by naive animals to the point of being considered demonstrator animals. Learning progress of naive rats in 40 trials, calculated as percentage of successful animals for each trial (n = 14). Error bars are mean ± standard error of the mean (SEM).
